# The conservation landscape of the human ribosomal RNA gene repeats

**DOI:** 10.1101/242602

**Authors:** Saumya Agrawal, Austen R.D. Ganley

**Affiliations:** Institute of Natural and Mathematical Sciences, Massey University, Private Bag 102-904, Auckland 0632, New Zealand; School of Biological Sciences, University of Auckland, Private Bag 92019, Auckland 1142, New Zealand (current address)

## Abstract

Ribosomal RNA gene repeats (rDNA) encode ribosomal RNA, a major component of ribosomes. Ribosome biogenesis is central to cellular metabolic regulation, and several diseases are associated with rDNA dysfunction, notably cancer, However, its highly repetitive nature has severely limited characterization of the elements responsible for rDNA function. Here we make use of phylogenetic footprinting to provide a comprehensive list of novel, potentially functional elements in the human rDNA. Complete rDNA sequences for six non-human primate species were constructed using *de novo* whole genome assemblies. These new sequences were used to determine the conservation profile of the human rDNA, revealing 49 conserved regions in the rDNA intergenic spacer (IGS). To provide insights into the potential roles of these conserved regions, the conservation profile was integrated with functional genomics datasets. We find two major zones that contain conserved elements characterised by enrichment of transcription-associated chromatin factors, and transcription. Conservation of some IGS transcripts in the apes underpins the potential functional significance of these transcripts and the elements controlling their expression. Our results characterize the conservation landscape of the human IGS, and suggest that noncoding transcription and chromatin elements are conserved and important features of this unique genomic region.

## Introduction

A characteristic feature of most eukaryote genomes is the presence of one or more tandem arrays of gene repeats encoding ribosomal RNA (rRNA), a key building block of ribosomes. The major eukaryotic rRNA gene repeat family is known as the ribosomal DNA (rDNA), with each repeat encompassing a coding region encoding 18S, 5.8S and 28S rRNA, and an intergenic spacer (IGS) that separates adjacent coding regions (**Fig 1**). In humans, each repeat unit is ~43 kb in length, with a ~13 kb rRNA coding region and a ~30 kb IGS [1], and there are approximately 200-600 rDNA copies distributed amongst tandem arrays on the short arms of the five acrocentric chromosomes (chromosomes 13, 14, 15, 21, and 22) [2–6]. The rDNA is transcribed by RNA Polymerase I (Pol-I) in the nucleolus [7,8], and this primary role in ribosome biogenesis places the rDNA at the heart of cellular metabolic homeostasis [9]. In addition, the rDNA has been found to mediate a number of “extra-coding” functions, including roles in genome stability [10,11], cell cycle control [12–16], protein sequestration [17], epigenetic silencing [18,19], and aging [20,21].

**Fig 1:**
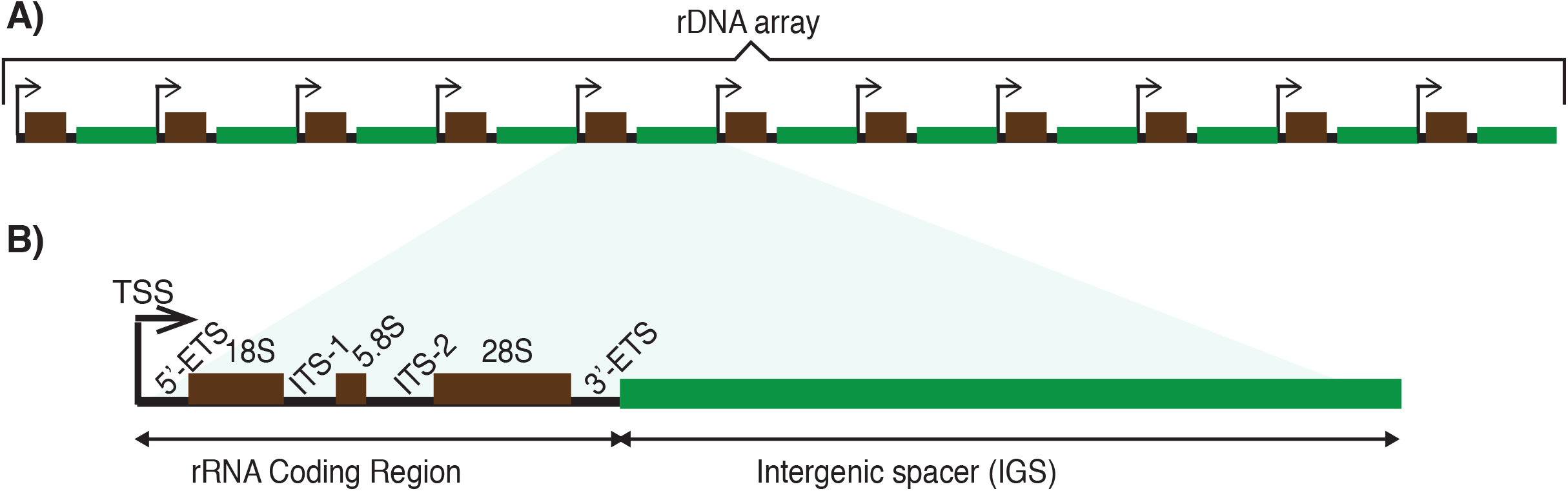
Eukaryotic ribosomal DNA organization. **A**) Head-to-tail tandem arrangement of rDNA repeat units. Typically there are more units in an array than depicted. **B**) Each rDNA unit has an rRNA coding region (black) and an intergenic spacer (IGS; green). The coding region encodes the ~18S, 5.8S and ~28S rRNAs (black boxes) separated by two internal transcribed spacers (ITS-1 and 2) and flanked by two external transcribed spacers (5’- and 3’-ETS).

A critical outcome of the central role the rDNA plays in the biology of the cell is an association with a number of human diseases. The best characterized is the association between ribosome biogenesis/rDNA and cancer, which dates back over 100 years and stems from observations of nucleolar hypertrophy and upregulated rRNA expression in tumour cells [22–25]. rRNA dysfunction is also associated with a group of genetic diseases that result from impaired ribosome biogenesis, known as ribosomopathies [26,27]. In addition, there is growing evidence for the rDNA playing a role in cellular differentiation [28–31]. Despite these strong connections to human pathology, the rDNA remains poorly characterized. Although an rDNA unit sequence is available in Genbank (accession U13369), it has not been placed in the human genome chromosomal assembly [32], and consequently the rDNA is excluded from most genome-wide analyses. Moreover, the lack of tools to genetically manipulate the highly repetitive rDNA in mammalian systems means that the human rDNA is not well characterized at the molecular level.

Work in *Saccharomyces cerevisiae* has shown that many rRNA regulatory and rDNA extra-coding functions are encoded in the IGS, where a number of functional elements have been characterized [10,33–46]. In stark contrast, even though the human IGS is approximately ten times longer than yeast, few functional elements have been defined to date. Those that have are restricted to the rRNA promoter [47], 10 bp repeats (Sal boxes) some of which act as terminators of the primary rRNA transcript [48], and two noncoding IGS transcripts that are associated with stress response [17]. Other elements have been identified from their sequence composition, including several other repeat elements [1], a cdc27 pseudogene [49], and putative c-Myc and p53 binding sites [50,51]. Pioneering work characterizing the chromatin structure of the human IGS has provided evidence for regions with distinct chromatin states, including states characteristic of transcriptional regulatory activity [52]. Furthermore, there appears to be dynamic regulation of this rDNA chromatin structure [53,54]. However, without further characterization, the functional significance of these human rDNA chromatin elements is unclear.

Comparative genomics is a powerful method for the identification of functional elements that are difficult to detect by traditional molecular approaches [55–58]. In particular, phylogenetic footprinting is an effective way to identify potentially functional elements using orthologous sequence data alone. The principle is that mutations in functional elements will be deleterious, therefore changes in the sequences of functional elements are selected against and change at a slower rate than non-functional elements over evolutionary time [59]. Thus, comparison of orthologous sequences from related species results in the functional elements appearing as “phylogenetic footprints” - highly conserved regions in a multiple sequence alignment against a background of non-functional, poorly conserved sequences [59]. Application of this method to the rDNA of *S. cerevisiae* successfully identified both known and novel functional elements in the IGS [10,44]. Given how little is known about functional elements in the human IGS and the strong connections between rDNA biology and human pathology, we decided to utilize phylogenetic footprinting to identify potential functional elements in the human rDNA.

Here, we constructed complete rDNA sequences from six primate species for which these sequences were previously unknown. Alignment of these sequences with a human rDNA sequence shows that previously identified functional elements in the human IGS are evident as phylogenetic footprints, and there are a number of other conserved regions not associated with any known functional element. Building on the results characterizing the chromatin state of the human rDNA [52], we shed light on the potential functions of these uncharacterized IGS conserved regions by overlaying publicly available RNA-seq, CAGE, and ChIP-seq data onto the conservation profiles. These analyses suggest that chromatin structure and the production/regulation of noncoding transcripts are major activities associated with sequence conservation in the human IGS. This is reinforced by conservation of IGS transcriptional activity in the apes, implying that these activities may be important for human rDNA function.

## Materials and Methods

### Whole genome assemblies to obtain the primate rDNA sequences

Whole genome sequencing (WGS) data for the six primates *viz*. chimpanzee (*Pan troglodytes*), gorilla (*Gorilla gorilla*), orangutan (*Pongo abelii*), gibbon (*Nomascus leucogenys*), rhesus macaque (*Macaca mulatta*), and common marmoset (*Callithrix jacchus)* were obtained from the Ensemble database (**S1 Table**). Whole genome assemblies (WGAs) for chimpanzee, gorilla, gibbon, macaque and common marmoset were performed using Arachne ver. r37405, and orangutan using Arachne ver. r37578, on a 64-bit server with six-core an Intel Xeon @ 2.67GHz processor and 512 GB RAM. We used Arachne [60,61; Supplemental Table S1-S2], as it resolved the rDNA unit the best in a comparative study of whole genome assemblers that we performed [62]. Default parameters were used for all assemblies. The steps to construct complete rDNA sequences are given in **S1 Supporting Methods Section 1.1**.

### BAC filters screening and BAC clones

BAC filters and *E. coli* containing the rDNA BAC clones were obtained from Children’s Hospital Oakland Research Institute, USA (CHORI; http://www.chori.org) (**S3 Table**). A 594 bp human 18S rDNA PCR product probe (Genbank U13369 coordinates 4,328-4,922) was made using male human template genomic DNA (Promega), primers HS_18S_rDNA_F (5′-AGCTCGTAGTTGGATCTTGG- 3′) and HS_18S_rDNA_R (5′- GTGAGGTTTCCCGTGTTGAG –3’), and DIG high prime DNA Labeling Kit II (Roche). To identify rDNA-containing BAC clones, Southern hybridization was used to screen the BAC filters with chemiluminescent detection and CDP-Star (Roche). BAC extraction was performed using overnight LB/chloramphenicol (30 μg/L) *E. coli* cultures containing the BAC of interest with the NucleoBond® Xtra Maxi Plus (Macherey-Nagel) kit.

### Determination of the primate rDNA size

To determine rDNA unit size, 10 μl of purified BAC DNA was digested with 100U of I-P*po*I (Promega) overnight. I-P*po*I digested products were run on 1% pulsed field certified agarose (BioRad) in 0.5X TBE gels with a CHEF Mapper XA (Bio-Rad) for 31 hrs using FIGE settings 180 V and 120 V forward and reverse voltages, respectively, and a 0.4 sec to 2 sec linear ramp switch time at 14°C. To aid resolution of bands, 5 kb ladder (Bio-Rad) was mixed with loading dye and water in a 1:1:2 ratio and incubated for 2 hrs at 37°C, 50°C for 15 min, and on ice for 10 min.

### rDNA BAC sequencing and analysis

Indexed libraries were prepared from BAC clones using NucleoBond® Xtra Maxi Plus (Macherey-Nagel). NGS was performed using Illumina HiSeq 2000 with 2×100 bp paired end reads and a 250 bp insert size. Low quality (<13) ends of reads were trimmed off and reads <25 bp in length were removed using SolexaQA [63]. Processed reads were mapped to the corresponding WGA rDNA sequence using bowtie (ver. 0.12.8). Consensus sequences were generated using a minimum coverage cutoff of 5 with CLC Genomic workbench and aligned to the corresponding WGA rDNA sequence using the MAFFT server [http://mafft.cbrc.jp/alignment/server; 64] with strategy E-INS-I and scoring matrix 1 PAM. Repeat regions in the rDNA sequences were identified using RepeatMasker (http://www.repeatmasker.org) with “DNA source” set as “human”. Alu elements in the IGS were confirmed using DFAM database (ver. 1.1) [http://dfam.janelia.org; 65], and numbered according to their IGS position (starting closest to the 3’-ETS). Other sequence elements in the IGS were identified using YASS [66] and BLAST [67].

### Multiple sequence alignment and similarity plots

Primate rDNA sequences were aligned to the human rDNA sequence (**S1 Appendix**) to generate multiple sequence alignments (MSA) using MAFFT (ver. 6.935b) [64,68] with strategy E-INS-i (-- genafpair), 1 PAM scoring matrix (--kimura 1), and gap penalty zero (--ep 0) (command: mafft -- genafpair --maxiterate 6 --thread 6 --cluastalout --kimura 1 --ep 0 --reorder fasta_input_file > seq.aln). Where required, alignments were adjusted by visual inspection. Columns with gaps in the human rDNA reference sequence were removed before similarity plot construction using Synplot [http://hscl.cimr.cam.ac.uk/syn_plot.html; 69] with a sliding window of 50 and increments of 1 bp. Human rDNA annotations were mapped onto the similarity plot using GFF files.

### Identification of conserved regions

Conserved regions in the MSA were identified using phastCons [70,71] using the phylogeny matrix for 99 vertebrates obtained by ENCODE (http://hgdownload.cse.ucsc.edu/goldenPath/hg38/phastCons100way/hg38.phastCons100way.mod).

### ORC mapping and peak analysis

Single end reads (1×36 bp) for ORC ChIP-seq and corresponding Input [72] data were processed and mapped to the modified human genome assembly using bowtie ver. 0.12.8 (parameters: -l 30 -n 3 -a --best --strata -m 1). Mapped reads were sorted and duplicate reads were removed using Picard (**S1 Supporting Methods Section 1.2**). ORC enrichment was determined and noise removed using MACS2 (**S1 Supporting Methods Section 1.2**). MACS2 function bdgpeakcall [pvalue cutoff 10^−20^ (-c 20)] was used to identify ORC peaks, and enrichment and peaks were visualized using Integrative Genomics Viewer (IGV) ver. 2.3.

### Transcriptome profiling

We introduced the human rDNA sequence into chr21 of the human genome assembly (hg19) to produce a modified human genome assembly. Long RNA-seq [poly(A+) and poly(A−)] and small RNA-seq data for human cell lines HUVEC, GM12878, H-1hESC, K562, HepG-2, and HeLa-S3 were obtained from the CSHL long RNA-seq and short RNA-seq databases, respectively (**S2 Appendix**). The long RNA-seq data were mapped to the modified human genome assembly using STAR aligner (ver. 2.2.0) [73]. Mapped reads were assembled (reads mapping to the rDNA coding region were masked) using Cufflink (ver. 2.2.1) [74]. Details are given in **S1 Supporting Methods Section 1.3**. Small RNA-seq data were mapped to the modified human genome assembly using bowtie. Regions with >= 5 read coverage were extracted using bedtools.

CAGE data for human cell lines HUVEC, GM12878, H9-hESC, K562, HepG-2, and HeLa-S3 were obtained from FANTOM [75; Supplemental data 3], and mapped to the modified repeat masked human genome assembly using bowtie (ver. 0.12.8). Masked assemblies were used to avoid multimapping reads from repeat regions, as CAGE reads are single end 35 bp reads with pseudo quality values. Paraclu (ver. 3) was used to identify the tag enrichments. Details are given in **S1 Supporting Methods Section 1.4**.

Paired-end (2×101 bp) total RNA-seq data for heart, kidney, liver, lung, and skeletal muscle of chimpanzee were obtained from the Nonhuman Primate Reference Transcriptome Resource [76], and analyzed as for the human RNA-seq analysis (**S1 Supporting Methods Section 1.3**). Poly(A+) single end data (1×76 bp) from heart, kidney, and liver of orangutan and macaque were obtained from Brawand *et. al*. [77], and analyzed as for the human RNA-seq analysis (**S1 Supporting Methods Section 1.3**) except that the STAR aligner parameter “outFilterMismatchNmax” was set to “5” and the Cufflink parameter “library-type” was change to “fr-unstranded”.

### Chromatin profiling

Data for histone modifications (H3K4Me1, H3K4Me2, H3K4Me3, H3K9Ac, H3K27Ac, H2.AZ, H3K36Me3, H3K9Me1, H4K20Me1, H3K79Me2, H3K27Me3 and H3K9Me3), RNA polymerases (Pol II and Pol III), transcription factors (TBP, ZNF143, c-Myc, Brf3, Brf1, Brd1, and UBF), CTCF, and Input for cell types HUVEC, GM12878, H-1hESC, K562, HepG-2, HeLa-S3, and A549 were downloaded from ENCODE [78; S2 Appendix]. Reads were processed and mapped to the modified human genome assembly using bowtie (ver. 0.12.8). Enrichment peaks were called using macs2 [79]. Details are given in **S1 Supporting Methods Section 1.2**. Mapped chromatin markers were combined to predict the rDNA chromatin states using Segway, for which a 10 state model underwent unsupervised training on 1% of the human genome [80] before prediction of chromatin states.

### Availability of data and material

The human rDNA sequence from BAC clone GL000220.1 is available (**S1 Appendix**). Primate rDNA sequences constructed using WGS data and sequencing of BAC clones are available from Genbank (accessions KX061886-KX061891 and KX061874-KX061885, respectively). Raw NGS data for primate rDNA BACs are available from the Sequence Read Archive (accession SRP068821). IGV sessions for visualizing the rDNA ChIP-seq peaks, RNA-seq predicted transcripts and CAGE peaks for cell types included in this study are available through figshare (**https://doi.org/10.17608/k6.auckland.6159395.v1**).

## Results

### Selection of species for phylogenetic footprinting

We set out to use phylogenetic footprinting to identify regions in the human IGS that are potential functional but have escaped detection because of the difficulties of working with the highly repetitive rDNA region. To do this, we decided to compare the human rDNA sequence with rDNA sequences from various primates. However, despite the genomes of several primate species having been sequenced, the complete rDNA sequence has not been identified, therefore we constructed rDNA sequences for selected primate species using whole genome assemblies (WGA). In our hands short-read next generation sequencing data are refractory to the assembly of complete rDNA units (unpublished results). Therefore, the first criterion for choosing the primate species for sequence comparisons was the availability of Sanger WGS data. The range of species relatedness is critical for phylogenetic footprinting [81], therefore our second criterion was inclusion of species with varying relatedness to human. Based on these criteria, we selected six primates (of the roughly 300 living species of primates distributed among 13 families [82]) that had Sanger whole genome sequence data available [83]: *Pan troglodytes* (chimpanzee), *Gorilla gorilla* (gorilla), and *Pongo abelii* (orangutan) from the Hominidae, *Nomascus leucogenys* (gibbon) from the Hylobatidae, *Macaca mulatta* (rhesus macaque) from the old world monkeys, and *Callithrix jacchus* (common marmoset) from the new world monkeys. These primates include both species closely related to human (Hominidae and Hylobatidae), together with more distantly related species (old and new world monkeys) (**Fig 2A**).

**Fig 2:**
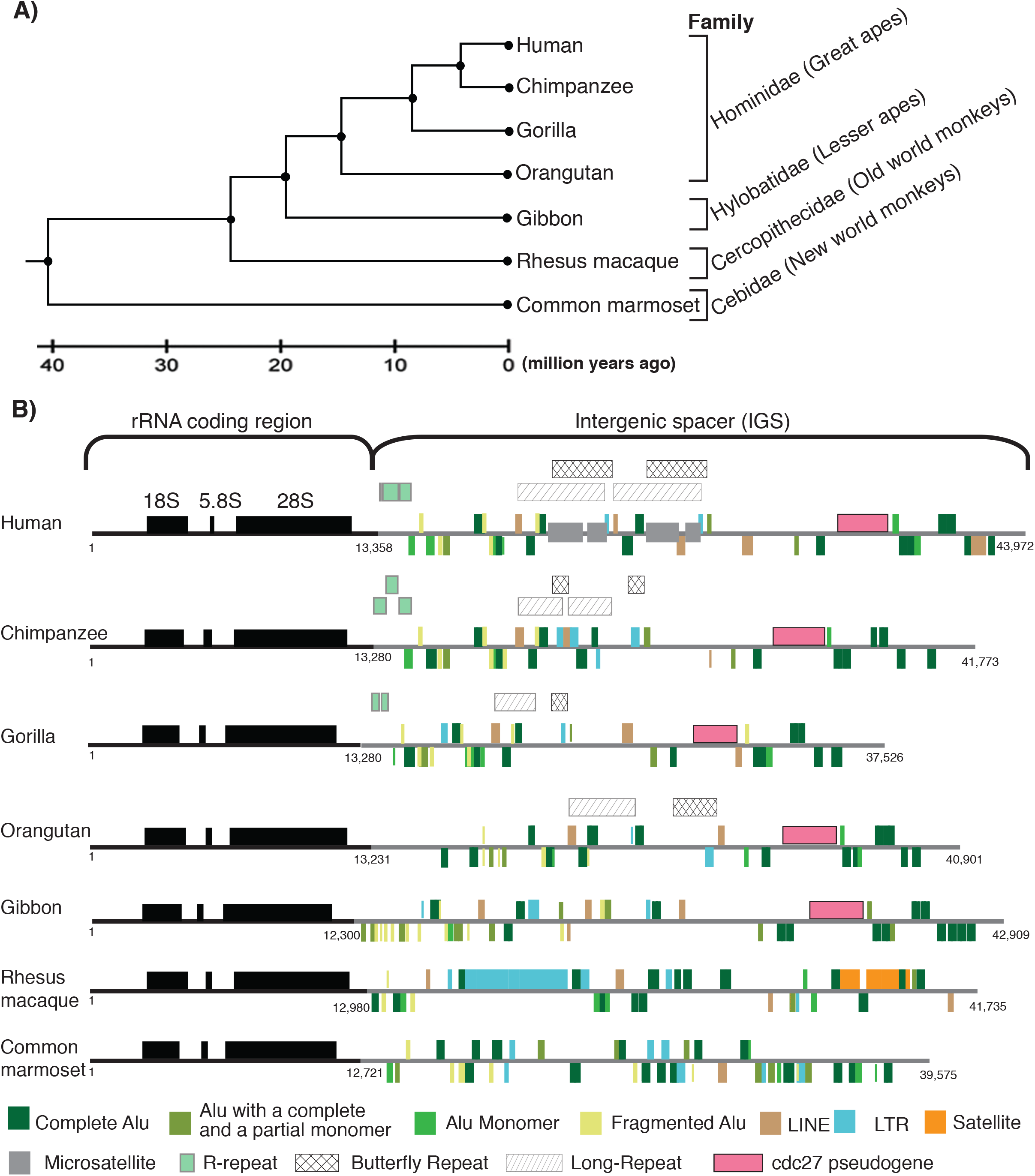
Primate rDNA repeat units. **A**) Phylogenetic tree showing the relationships between primate species selected for rDNA phylogenetic footprinting [adapted from 84]. **B**) Human and primate rDNA unit structures are shown. The rRNA coding region (black line), including the 18S, 5.8S and 28S rRNA subunits (black boxes), and the IGS (grey line) are indicated along with the positions of repeat elements and a cdc27 pseudogene. Elements above the line are on the forward strand; those below on the reverse strand. The rRNA coding region/IGS coordinates and rDNA unit lengths are indicated.

### Reference human rDNA sequence

The widely used reference human rDNA unit (Genbank accession U13369) was constructed by assembling several partial sequences obtained by different labs [1]. This sequence is known to contain errors [22,85], hence we wanted to use a human rDNA sequence from a single source that is likely to have fewer errors. We chose the complete human rDNA unit sequence (43,972 bp) present in an unannotated BAC clone (Genbank accession GL000220.1; **S1 Appendix**) [32] that contains a complete and partial rDNA unit together with a part of the rDNA distal flanking region [32]. We refer to this complete single rDNA sequence we use as the “human rDNA”, and it includes a 13,357 bp coding region and a 30,615 bp IGS (as determined by comparison to the Genbank human rDNA sequence). Excluding copy number variation in microsatellite and other repeats in the IGS (**S3 Appendix Section 1.1**), the human rDNA shows 98.1% sequence identity to U13369.

### Constructing primate rDNA sequences

To perform phylogenetic footprinting, we first constructed rDNA sequences for the selected primate species using WGA. The high level of sequence identity between rDNA units within a genome [86–88] leads genome assemblers to construct a single, high-coverage “consensus” rDNA unit sequence from the multiple rDNA repeats. The coverage level will be greater than that of unique regions by a factor of the rDNA copy number (about 200-500 in primates; [89,90]). We therefore performed WGA on publicly available WGS data for the primate species (**S1-S2 Tables**) and selected high-coverage contigs. These contigs were screened using the human rDNA sequence to identify rDNA-containing contigs, and merged to produce complete rDNA sequences. From this we obtained rDNA units for the six primate species, ranging in size from 37.5 – 42.9 kb (**Fig 2B**), and the regions corresponding to the rRNA coding region and IGS were identified by comparison with the human rDNA (**S4 Table**). The human coding region aligns completely (end to end) to all primate rDNA sequences except marmoset, for which the 5’ ETS is 272 bp shorter than the human 5’ ETS. This may be because the marmoset 5’ ETS is actually shorter than human, or because the WGA failed to properly assemble this region.

Use of the human rDNA to identify rDNA contigs in the primate WGAs makes it possible that regions present in other primates, but not in human, were missed. Furthermore, the presence of repetitive elements in the IGS that are also found in other regions of the genome [91] may have led to WGA errors [92]. To eliminate these possibilities, we first identified rDNA-containing BAC clones for the primate species (except chimpanzee, which has a high level of genomic sequence identity to human) by screening BAC genomic libraries (**S3 Table**). We compared the sizes of the WGA and BAC rDNA units by digesting the BAC clones with I-*Ppo*I, a homing enzyme that cuts only once in the rDNA (in the 28S), separating the fragments using field inversion gel electrophoresis (FIGE), and performing Southern hybridization (**S1 Fig**). The estimated lengths of the BAC (via FIGE) and the WGA rDNA sequences are similar (**S1 Fig; S5 Table**), with the FIGE sizes being consistently ~1 kb larger than the WGA sizes (**S5 Table**). We favor the interpretation that the FIGE gels slightly overestimate the size, given the consistent differences in size and the end-to-end matching to the human rDNA sequence. However, we cannot rule out the primate species all contain an additional ~1 kb region that is not present in human and does not assemble. Nevertheless, the results suggest that the primate rDNA units we assembled contain the sequence information necessary to determine the conservation profile of the human rDNA. To further confirm the integrity of the WGA rDNA sequences, the primate rDNA BAC clones were sequenced, and consensus primate rDNA sequences were obtained by mapping the reads to the corresponding WGA rDNA sequences. On average, the consensus BAC rDNA sequences are >97% identical to the WGA sequences (**S6 Table**). The variation is mainly due to gaps in the rRNA coding regions caused by an absence of reads from these regions in the NGS data, a phenomenon we have observed previously (unpublished results). The high level of sequence identity (where reads are present) suggests the WGS rDNA sequences are accurate representations of the true rDNA sequences. Therefore, we used the WGA sequences as the reference rDNA sequences for all non-human primate species.

Next, we characterized these new primate rDNA sequences to determine their structural similarity to the human rDNA. The length of the coding region in the six primate species is similar to human *i.e*. approximately 13 kb, except gibbon that is slightly smaller (**S4 Table**). As expected, as we move from chimpanzee to common marmoset, the pairwise sequence identity with human decreases for the coding region (**S4 Table**). The microsatellite component of the rDNA unit in all six primate species is higher than the genome wide average for each species **Table 1**), and human has the highest microsatellite content because of two long, unique [TC]_n_ repeat blocks (**Fig 2B**). Alu elements are the most abundant repeat element in the primate IGS (**Table 1**), and a number are orthologous between human, apes and rhesus macaque (**S2 Fig; S7 Table; S3 Appendix Section 1.2**). We found that, consistent with a previous report [49], Alu_human_22, Alu_human_25 and Alu_human_27 are present in chimpanzee, gorilla, orangutan, gibbon, and rhesus macaque, while Alu_human_23 is present in apes but not rhesus macaque. It has also been reported that orthologs of Alu_human_26 and Alu_human_28 are present in rhesus macaque [49], but our results show that while these two Alus are conserved in apes, the Alu elements present in similar regions in rhesus macaque are on the opposite strand. Several repeats of unknown function have been identified in the human rDNA (called Long repeats and Butterfly repeats; [1]). These show varying distributions amongst the primates (**Fig 2B**), suggesting they originated at different points in primate evolution. The pseudogene of cdc27 in the human IGS is also present in apes but not in monkeys, as previously reported [49], and the rhesus macaque rDNA sequence contains large LTR retrotransposons and satellite repeats that are absent from the other species (**Fig 2B**). Overall, these results show that a clear signal of orthology and synteny is retained in the rDNA sequences of the selected primates, but there is also sufficient diversity for phylogenetic footprinting to be effective.

**Table 1:**
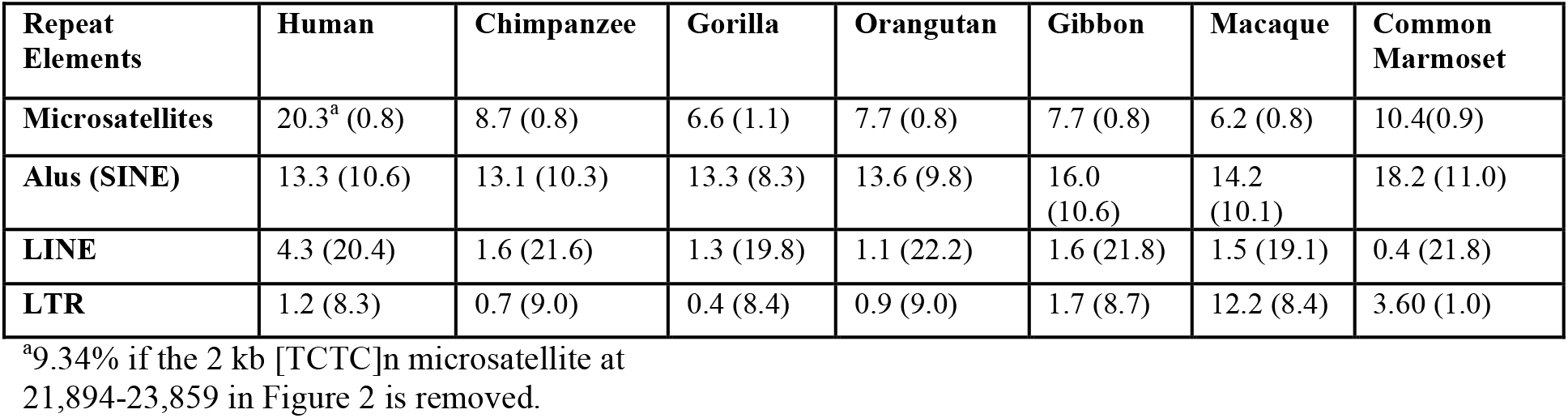
Repeat composition of the primate rDNA sequences as a percent of total rDNA length (with genome-wide percent abundance in parentheses for comparison).

| Repeat Elements | Human | Chimpanzee | Gorilla | Orangutan | Gibbon | Macaque | Common Marmoset |
| --- | --- | --- | --- | --- | --- | --- | --- |
| **Microsatellites** | 20.3<sup>a</sup> (0.8) | 8.7 (0.8) | 6.6 (1.1) | 7.7 (0.8) | 7.7 (0.8) | 6.2 (0.8) | 10.4(0.9) |
| **Alus (SINE)** | 13.3 (10.6) | 13.1 (10.3) | 13.3 (8.3) | 13.6 (9.8) | 16.0 (10.6) | 14.2 (10.1) | 18.2 (11.0) |
| **LINE** | 4.3 (20.4) | 1.6 (21.6) | 1.3 (19.8) | 1.1 (22.2) | 1.6 (21.8) | 1.5 (19.1) | 0.4 (21.8) |
| **LTR** | 1.2 (8.3) | 0.7 (9.0) | 0.4 (8.4) | 0.9 (9.0) | 1.7 (8.7) | 12.2 (8.4) | 3.60 (1.0) |

<sup>a</sup>9.34% if the 2 kb [TCTC]<sub>n</sub> microsatellite at 21,894-23,859 in Figure 2 is removed.

### Conserved regions in the human IGS identified by phylogenetic footprinting

To identify novel conserved regions that are potentially functional in the human rDNA through phylogenetic footprinting, we aligned the human and primate rDNA sequences. Although the human and common marmoset rDNA sequences align, the alignment is compromised by the relatively low level of sequence identity (**S4 Table**). Therefore, an alignment with the common marmoset omitted (MSA_human-macaque_) was used for the phylogenetic footprinting. The MSA_human-macaque_ has long runs of gaps that are predominantly the result of satellite blocks in the rhesus macaque rDNA. Because the goal was to identify conserved regions in the human rDNA, all columns in the MSA with gaps in the human rDNA were removed. To observe the level of sequence conservation, a similarity plot was generated using Synplot (**Fig 3**). We then identified the regions that are conserved using phastCons, which employs maximum likelihood to fit a phylogenetic hidden Markov model to the alignment [70]. Forty-nine conserved regions (c-1 to c-49) were identified in the human IGS (**Fig 3; S8 Table**), corresponding to 21.9% of its length. These conserved regions map to both unique regions and Alu elements in the rDNA (**Fig 3**). We looked to see if these regions are also conserved in the common marmoset and mouse rDNA (using Genbank rDNA reference accession BK000964.3). Twenty-three conserved regions mapped to the common marmoset rDNA, and four mapped to the mouse rDNA, with three found in both, using a >50% identity threshold (**Fig 3; S9 Table**). Interestingly, two of the three regions conserved with both mouse and common marmoset (c35-36) cover a single Alu repeat (Alu_human_20) with no described function. Together, this phylogenetic footprinting approach reveals conserved regions in the human IGS, including some deeply conserved regions, that represent potentially functional elements.

**Fig 3:**
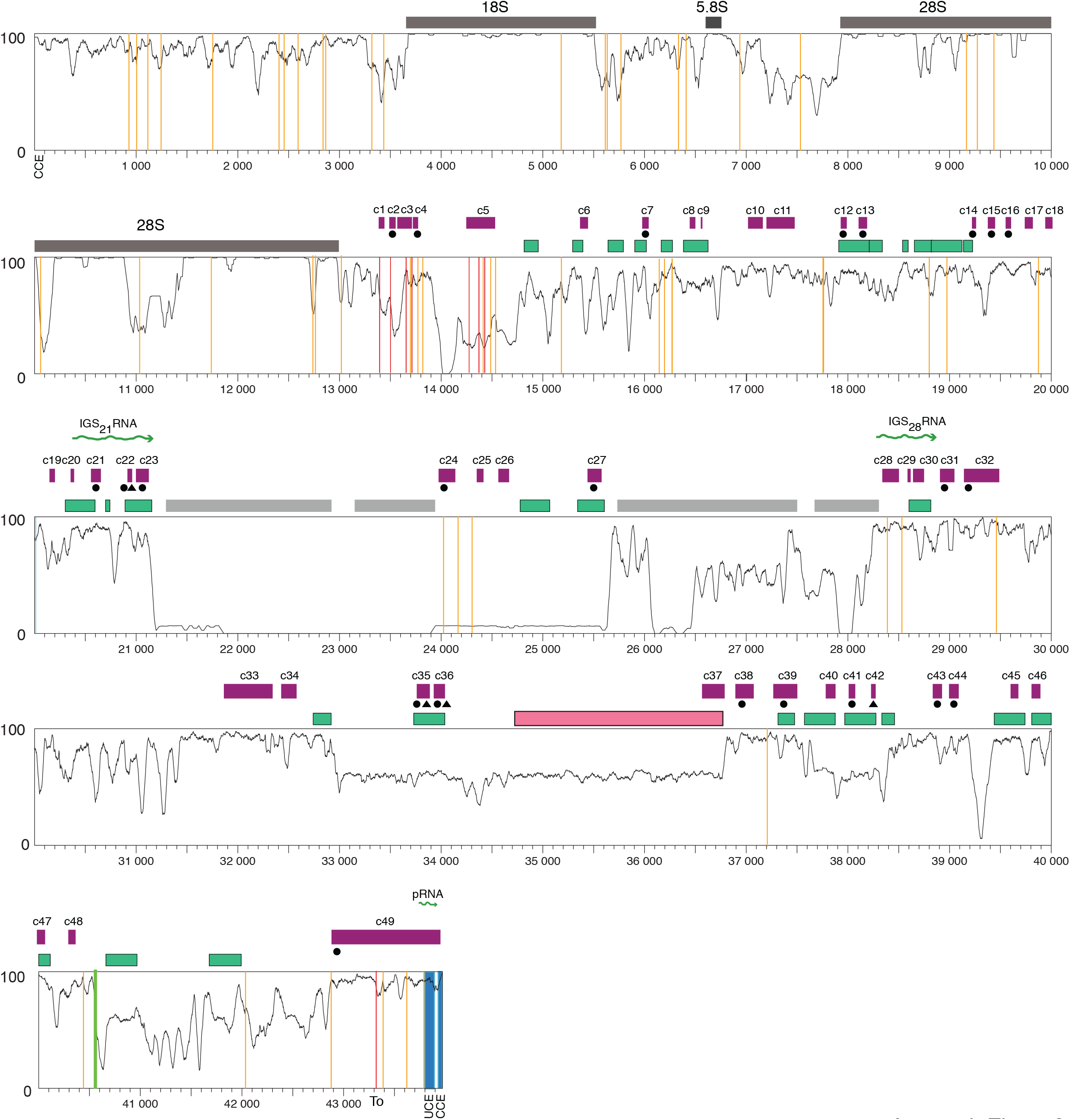
Sequence similarity plot of the primate rDNA. The horizontal axis represents the position in the human rDNA; the vertical axis the level of sequence similarity between 0 (no identity) and 1 (all bases the same). A 50 bp sliding window with 1 bp increment was used to generate the similarity plot. Conserved regions in the IGS (purple boxes) were identified using phastCons. The positions of Alu elements (green boxes), microsatellites (grey boxes), a cdc27 pseudogene (pink box), the rRNA promoter (blue lines), previously identified IGS noncoding transcripts (green wiggly lines), c-Myc binding sites (orange lines), p53 binding site (green line), and Sal boxes (terminator elements; red lines) are indicated. Conserved regions with a black circle or triangle below are conserved in common marmoset and mouse rDNA, respectively.

### Conservation of previously known features in the human IGS

To verify that the phylogenetic footprinting is capable of identifying functional elements in the human rDNA, we looked at whether known human rDNA elements are conserved amongst the primates. As anticipated, the 18S and 5.8S rRNA coding regions are highly conserved across the primates, while the 28S rRNA coding region consists of conserved blocks interspersed with variable regions, as previously reported (**Fig 3**) [93; Figure 3,94,95]. The rRNA promoter has two characterized elements: an upstream control element (UCE) from position −156 to −107 and a core control element (CCE) from position −45 to +18 [47], and both elements are conserved (**Fig 3; S3.A Fig**). Several potential rRNA transcriptional terminators (Sal boxes) are present downstream of the 28S rRNA coding region [48,96], and all are conserved (**S3.B Fig**). In addition, the Sal box proximal to the rRNA promoter [48] is conserved, although the functional significance of a terminator in this position is not clear. The c-Myc binding sites identified around the rRNA promoter fall in a conserved region (c49; **Fig 3**), with this area having been shown to bind c-Myc [50]. Several other predicted c-Myc binding sites in the IGS also fall into conserved regions, although the majority (including sites near the terminator that were shown to bind c-Myc) do not (**Fig 3**) [50]. However, conservation of the actual binding motif itself does not automatically translate to a conserved region because of the thresholds used to define conserved blocks (**S4 Fig**), and some c-Myc binding motifs around the terminator that are not in a conserved region are, nevertheless, conserved. The region corresponding to the pRNA, a noncoding RNA transcript that plays a role in rDNA silencing in mouse [97], coincides with conserved region c49, although it is not conserved with mouse (Fig 3). Two human IGS transcripts that are produced as a result of stress [called IGS21RNA and IGS28RNA; 17]) overlap conserved regions c20-c23 and c28-c30, respectively (**Fig 3**). The conservation of these noncoding IGS transcripts suggests that their function in stress response may be conserved in primates. Together, our results show that a number of elements in the rDNA that are known or have been suggested to be functional appear as conserved peaks, suggesting that our phylogenetic footprinting approach has the ability to identify functional elements in the IGS.

### Association of unknown conserved regions with transcription

Previously known functional elements account for 11 (c1-c3, c20-c23, c28-c30 and c49) of the identified 49 conserved regions. The remaining conserved regions remain uncharacterized, and these regions may represent novel functional elements. Therefore, we next looked for potential functions of these novel conserved regions. The presence of characterized noncoding transcripts in the human IGS [17,97,98], as well as their prominence in the rDNA of other organisms [10,99–101], led us to explore whether some of the conserved regions are associated with noncoding transcription. We mapped publicly available long poly(A+) and poly(A−) (>200bp), and small RNA (< 200 bp) RNA-seq data [102] from all six cell lines of the first two tiers of the ENCODE project to a modified human genome assembly to which we added the human rDNA sequence (“modified human genome assembly”), without repeats masked. The cell lines included two normal cell lines (HUVEC and GM12878), one embryonic stem cell line (H1-hESC), and three cancer cell lines (K562, HeLa-S3, and HepG-2). Several novel poly(A+) and poly(A−) transcripts were identified, including transcripts in common across all cell lines, and transcripts restricted to a subset of cell lines (**S5 Fig; S10-S21 Tables**). To identify potential transcriptional start sites (TSS) for these noncoding transcripts, we mapped CAGE data from the FANTOM5 project [75] to the modified human genome assembly with repeats masked (to prevent spurious alignment of the short CAGE sequence reads). Several CAGE peaks were identified that support the presence of some of the novel IGS transcripts (**S5 Fig; S22 Table**).

The presence of transcripts that originate from the human IGS implies that transcriptional regulators (e.g. promoters, enhancers and insulators) are present in the IGS, and may correspond to some of the conserved regions. Therefore, we mapped publicly available ENCODE ChIP-seq data for histone modifications, RNA polymerase-II and III, transcription factors (TBP, c-Myc and ZNF143), and the insulator binding protein CTCF, a highly conserved protein that is involved in the three-dimensional organization of chromatin [103–105], to the modified human genome assembly. We used ChIP-seq data from the six cell lines that were subjected to RNA-seq analysis, as well as from an additional cancer cell line (A549) from tier-3 of the ENCODE project. Several peaks of enrichment for these factors were identified (**S6-S12 Figs**), with those associated with active transcription being distinct and sharp, while those associated with transcriptional repression are comparatively broad, as previously observed [52]. Cell line HeLa-S3 is an exception as the histone modifications peaks associated with active transcription are broad as well. The GM12878 cell line has fewer prominent histone modification peaks than the other cell lines, probably because of loss of a substantial number of ChIP-seq reads during the quality control step for this cell line (data not shown). We then integrated the histone modification, CTCF, and Pol-II profiles for all seven cell lines using Segway [106] to determine putative chromatin states in the IGS (**S13 Fig**). Finally, we intersected the RNA-seq, CAGE, and chromatin state datasets with the conserved regions to identify transcripts and chromatin states that are potentially functionally conserved. This analysis revealed three prominent zones in the IGS containing several conserved regions that either show evidence for active transcription or have chromatin states associated with transcription (**Fig 4**). Together, these zones account for 18 of the 38 unknown conserved regions, including 14 of the 23 regions conserved with the common marmoset. The first zone is located near the rRNA transcriptional terminator, and we call this zone-1. It encompasses conserved regions c6 to c23 (~14.8 kb − 21.1 kb) (**Fig 4**), and contains a number of both poly(A+) and poly(A−) transcripts common to all cell lines (**S5 Fig**), many of which appear to be spliced. There are a number of peaks of histone modifications that indicate chromatin states associated with transcription, most prominently in the H1-hESC and HepG2 cell lines. A number of the putative transcripts appear to originate upstream of this zone, in a region that is enriched for chromatin states associated with active transcription and with CAGE peaks, but does not show sequence conservation. Zone-1 also contains the previously identified IGS_21_RNA noncoding transcript (**Fig 3**).

**Fig 4:**
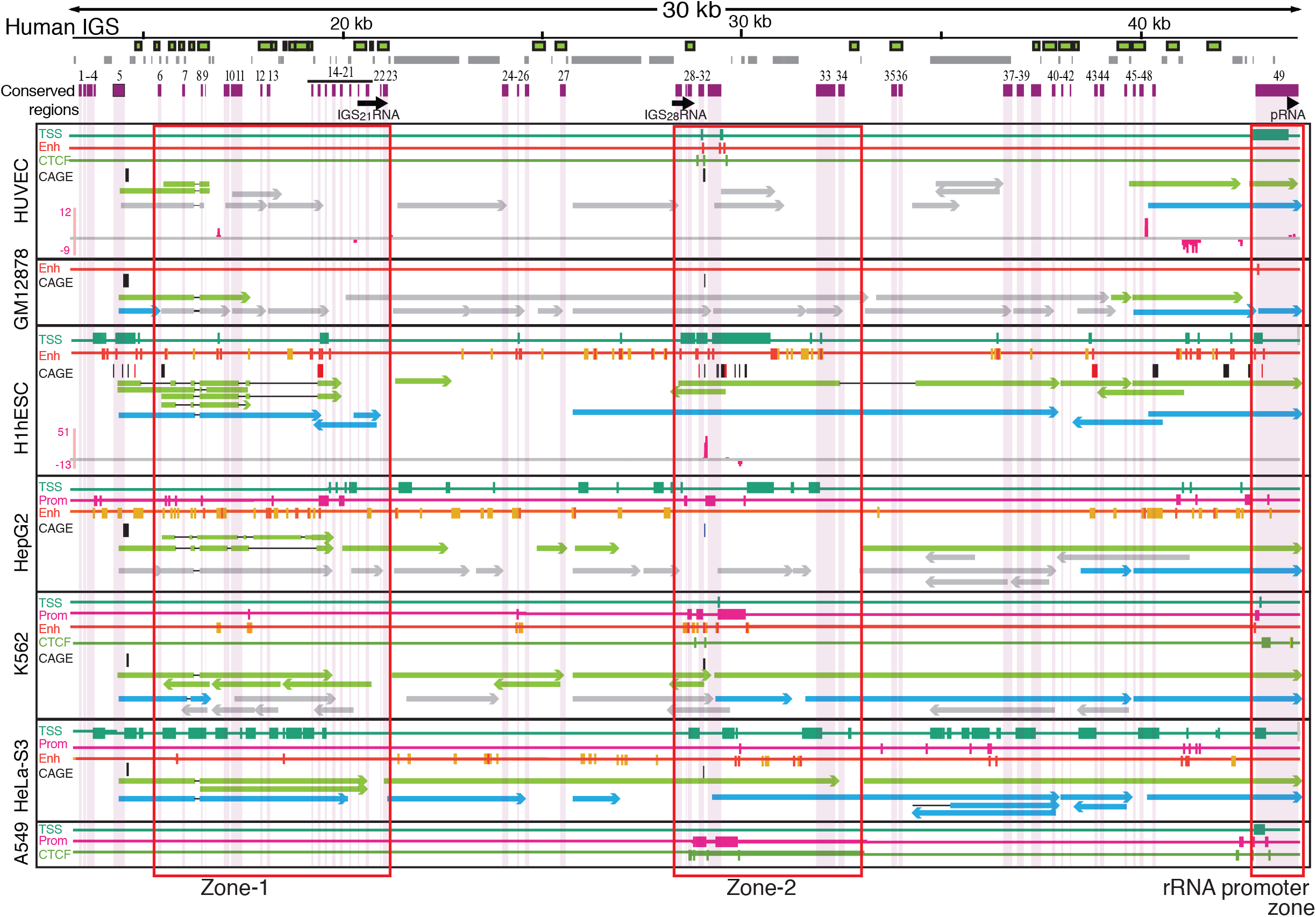
Two zones in the human IGS enriched for conserved regions and transcription associated factors. The human IGS is shown at top, with the positions of Alu elements (green boxes), microsatellites (grey boxes), conserved regions (purple boxes), and previously identified IGS noncoding transcripts (black arrows) indicated. Below are chromatin and transcriptional features of seven human cell lines. The positions of the conserved regions are indicated by pale shading. For each cell line the presence of transcriptional start site (TSS), promoter (Prom), enhancer (Enh), and CTCF segmentation states, obtained by merging histone modification, Pol II and CTCF peaks using Segway, are indicated. Below these, CAGE peaks are shown for the forward (black boxes) and reverse (red boxes) strands (CAGE stem cell data come from H9-hESC, not H1-hESC), followed by long poly(A+) and poly(A-) transcripts (green and blue arrows, respectively) with FPKM values >1; gray arrows indicate transcripts with FPKM < 1. Arrowheads indicate the direction of transcription. Peaks of small RNA are shown in pink. Zones 1 and 2 that are enriched for conserved regions and transcription-associated factors are boxed in red. Not all features have data available for all cell lines.

The second zone is roughly in the middle of the IGS, and we call this zone-2. It encompasses conserved regions c28-c34 (~28.2 to 32.6 kb; **Fig 4**) and shows strong enrichment for chromatin states associated with transcription and transcriptional regulation. Conserved regions c28-c30 correspond to the previously identified IGS_28_RNA noncoding transcript [17,52], and, consistent with previous results [52], show chromatin states associated with transcriptional activity (**Fig 4**). While we do not detect IGS_28_RNA specifically, we do find transcripts that overlap it. Conserved regions c31-c32 show an enrichment of active chromatin states, as reported previously [52], as well as transcripts in many cell lines (**Fig 4; S5 Fig**). This region also shows a peak of CAGE tags in the same position in all cell lines for which CAGE data are available (**Fig 4; S5 Fig**). Interestingly, there are two oppositely transcribed small RNA peaks in conserved region c31 that may represent transcription from a bidirectional promoter, and are only observed in H1-hESC (**Fig 4; S5 Fig**). In general, more CAGE tag peaks map in the stem cell line than the other cell lines, mirroring genome-wide patterns of embryonic stem cell expression [107] and suggesting the rDNA might be in an unusually permissive chromatin state for noncoding transcription in this cell type. Furthermore, zone-2 was the only part of the IGS for which CTCF segmentation states were predicted in all cell lines that had data.

The final zone encompasses the rRNA promoter (**Fig 4**). Noncoding transcripts are found in this zone (**S6-S11 Figs**), including small RNA peaks in the HUVEC cell line. Some of these transcripts may function like the mouse pRNA, a small RNA that influences rRNA transcription [97], with pRNA-like transcripts having been detected in the human rDNA before [52]. This zone also displays chromatin features characteristic of TSSs, promoters, and enhancers, depending on the cell line (**Fig 4**), and again, some of these features might relate to the presence of the pRNA. However, whether humans have a pRNA that is functionally equivalent to the mouse pRNA has not yet been determined.

Our analyses also show a number of poly(A+) and poly(A−) transcripts, small RNAs, and chromatin states associated with transcriptional activity outside of these zones. In some cases these overlap with conserved regions, but in other cases they do not, and it is difficult to determine whether the transcriptional features that overlap conserved regions are associated with the conservation or not. A number of the nonconserved transcriptional features correspond to microsatellite regions (**S12 Fig**), suggesting they might be artifacts of the spurious alignment of reads to IGS microsatellites [92]. However, microsatellites have been shown to act as promoters and/or enhancers [108–112], hence we cannot completely rule out that the chromatin states at these sites are real.

### Replication and double strand break association

The presence of origin of replication activity is a conserved feature of the rDNA [39,113–117]. Genome-wide mammalian origins of replication are not defined by sequence and there is not agreement on precisely where replication initiates in the rDNA [115,118–121]. We looked to see whether origin of replication complex association overlaps with conserved regions in case the rDNA initiates replication in a sequence-specific manner. We mapped origin of replication complex (ORC) ChIP-seq data [72] to the modified human genome assembly. The majority of ORC signal in the rDNA is found distributed across the rRNA coding region and the regions immediately flanking this (**Fig 5**). However, six smaller peaks of ORC enrichment are seen in the IGS, with five of them falling in conserved regions (**Fig 5**). These results suggest that the majority of replication in the human rDNA initiates in the rRNA coding region and/or the regions flanking it, consistent with reports that mammalian origins of replication are enriched in transcriptionally active regions [72]. Whether there is any biological significance to the minor ORC peaks at the conserved regions in the IGS is unclear.

**Fig 5:**
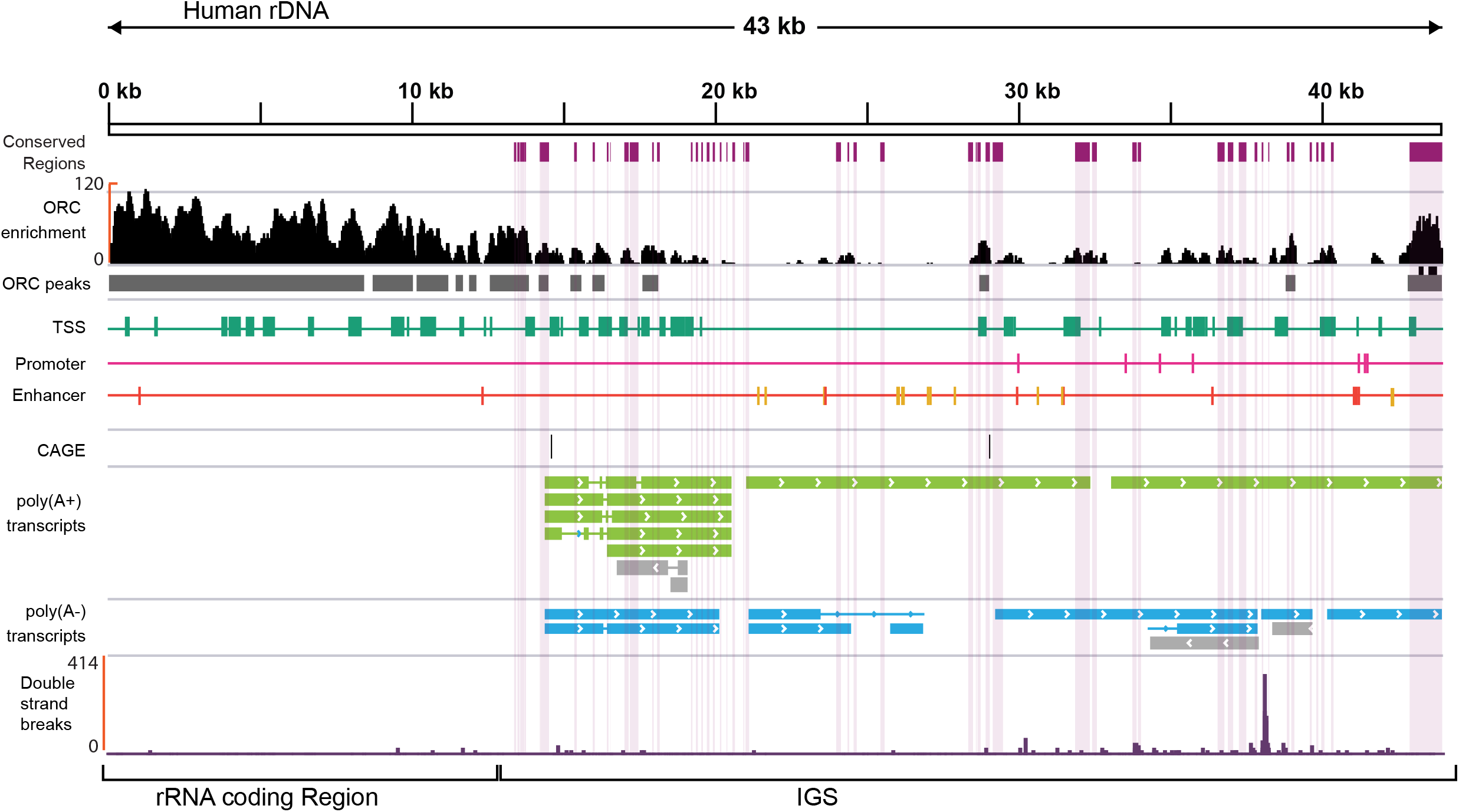
Origin replication complex (ORC) and double strand break (DSB) occurrence in the rDNA. The black plot represents enrichment of ORC in Hela-S3 cells and grey boxes below represent the position of peaks. Scale on the left is the -fold enrichment, and the scale above shows the position in the rDNA. Purple boxes represent conserved regions. The predicted chromatin states: transcription start site (TSS; green boxes), promoter (pink boxes), and enhancer (orange boxes) are shown. CAGE peaks are shown as black boxes (positive strand). Long poly(A+) and poly(A−) transcripts with FPKM values > 1 are shown as green and blue boxes, respectively. Gray arrows show transcripts with FPKM < 1. Arrows indicate the direction of transcription. The purple plot at bottom represents the DSB sites in HEK293T cells.

A key feature of the rDNA repeats in yeast is the presence of double strand breaks at a conserved site of unidirectional replication fork stalling known as the replication fork barrier site [42,43,122]. We examined whether recently reported double-strand break sites in the human rDNA [123] are located around conserved regions, but found no consistent pattern of association (**Fig 5**). Interestingly, however, the major DSB site in the rDNA lies in a region that is close to one peak of ORC enrichment, potentially suggesting the DSB site is a region of replication restart, such as observed at the yeast rDNA [124]. However, this site is at the opposite end of the IGS to where human replication fork barrier activity has been reported [125].

### Long noncoding RNAs are conserved among primates

Finally, we reasoned that the presence of transcripts and chromatin states associated with active transcription in conserved regions of the human IGS suggests that similar transcripts should be present in other primates. To test this, we took publicly available paired end total RNA-seq data from liver, lung, and skeletal muscle of chimpanzee [76], and single end poly(A+) RNA-seq data from liver, heart, and cerebellum of chimpanzee, orangutan, and macaque [77]. These data were mapped to the corresponding species’ genome assembly to which the appropriate rDNA sequence had been inserted. We found IGS transcripts in all tissues from chimpanzee and orangutan (**S14-S16Figs; S23-S26 Tables**), but in macaque such transcripts were only present in liver and heart tissue. We compared the primate IGS transcripts to HUVEC IGS transcripts, as HUVEC is a primary cell line that has a normal karyotype and is not artificially immortalized, hence is likely to be the closest to a “normal” human cell state. Transcripts similar to those found around the human promoter region are also found in chimpanzee and orangutan. In addition, transcripts similar to those found in zone-1 in the human IGS are found in all primate species we analyzed (**Fig 6**). Strikingly, there is conservation of splice junctions between human, chimpanzee and orangutan, even though the full lengths of the transcripts are not the same. No transcripts corresponding to zone-2 were found for the non-human primates analyzed here, and only one IGS transcript was found in macaque in zone-1, although this transcript does not overlap the HUVEC transcripts. Therefore, some but not all of the IGS transcripts that emanate from conserved regions in human are conserved across the apes, supporting the idea that these regions may have been conserved to maintain this transcriptional function. However, the lack of IGS transcripts in macaque suggests that transcriptional conservation does not extend as far as the monkeys, although we cannot rule out that the appropriate macaque tissues have not been sampled to find these IGS transcripts, or that their absence simply reflects a loss that is unique to macaque. The lack of transcripts from zone-2 in apes suggests that enrichment of transcriptional regulatory features in conserved regions in this zone may be involved with determining a specific chromatin structure, or that the production of transcripts is tissue-specific, such as the potentially stem cell-specific bidirectional RNA we identified in this region.

**Fig 6:**
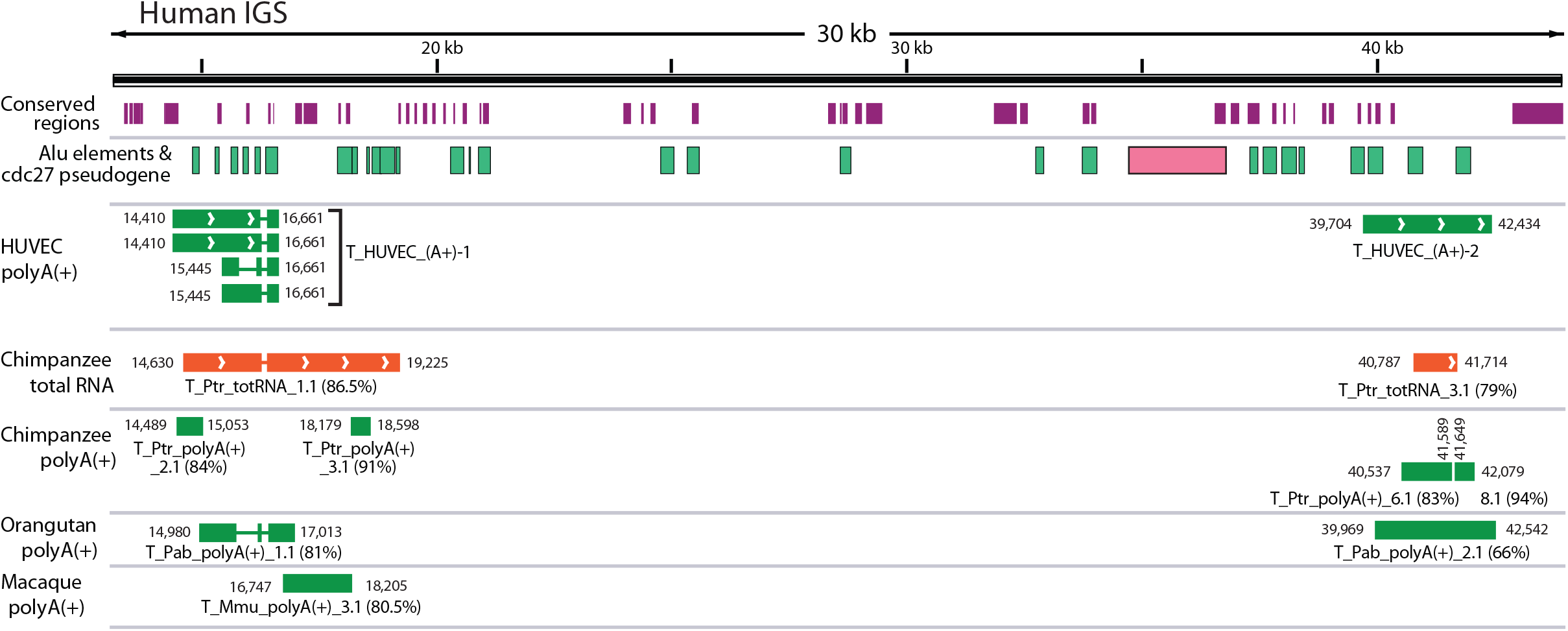
Conservation of human IGS transcripts amongst primates. The human IGS is indicated at top along with the conserved regions (purple boxes), Alu elements (green boxes) and cdc27 pseudogene (pink box). Below are poly(A+) IGS transcripts from the HUVEC cell line, followed by total RNA chimpanzee IGS transcripts (orange), and poly(A+) IGS transcripts from chimpanzee, orangutan, and rhesus macaque (green boxes). Only transcripts that are in common with human are shown. Transcript names and their start/end coordinates are indicated alongside, as are percent identities between each transcript and the human IGS (in parentheses). Arrowheads indicate direction of transcription.

## Discussion

In this study, we combined phylogenetic footprinting, a powerful tool to identify novel functional regions that are conserved over evolutionary time, with genomic datasets to overcome the challenges posed by the highly repetitive nature of the rDNA. In total, we identified 49 conserved regions in the human rDNA IGS. Several of these regions correspond to known functional elements, including the rRNA promoter and terminators, IGS noncoding transcripts, and protein binding sites, while others are novel. The novel conserved regions are dispersed throughout the IGS and correspond to both unique regions and repeat elements. The conserved regions identified here are restricted to elements that share a potential function with most of the primate species examined, and therefore do not include functional IGS elements that have evolved more recently in the lineage leading to humans. However, it may be possible to detect potential human-specific elements via determination of human accelerated regions [126]. Nevertheless, our results catalogue a large suite of potentially functional, uncharacterized regions in the human rDNA that will allow targeted investigations of their functionality. Our work has also provided complete rDNA reference sequences for six primate species that were previously unavailable. These new sequences will facilitate a better understanding of the rDNA in these primates and offer a strong comparative base for additional studies on the human rDNA.

Following the IGS chromatin state characterizations made by Scacheri and colleagues [52], we used several publicly available sequence databases to determine whether the conserved regions show distinctive chromatin states and/or noncoding transcripts that could provide evidence for the functions they putatively play. We found numerous long poly(A+) and poly(A−) transcripts in the human IGS, including many that have not been previously reported, suggesting there is pervasive transcription of the human IGS that is consistent with pervasive transcription in other regions of the genome [102,127,128]. Long noncoding RNAs from the IGS have been reported to be involved in regulating rRNA transcription [97] and stress response [17], therefore some of the novel long IGS transcripts we identified here may also be functional, and a number are conserved in part or whole. However, as for much of the genome-wide pervasive transcription, further work is required to determine what functions, if any, the novel IGS transcripts we document here have.

Mapping of chromatin datasets to the rDNA revealed several regions with chromatin structures that are consistent with transcriptional activity in the IGS, and with those previously reported [52]. Importantly, many of these putatively regulatory regions overlap conserved regions. In particular, two zones show a preponderance of conserved regions and features associated with transcription. Long poly(A+) transcription from zone-1 and near the promoter region is consistently observed in all the cell lines we examined, and some, but not all, of this transcriptional activity is reinforced by chromatin marks associated with active transcription. The presence of transcripts emanating upstream of zone-1 in all cell lines is striking (**Fig 4; S5 Fig**), although the exonic structure of these transcripts is variable and their expression in different cell lines is also variable (**S17 Fig**). While these could represent read-through rRNA transcription, there are three reasons to suggest they do not. First, they are present as both polyA+ and polyA-transcripts, whereas if they were read-through rRNA transcripts, polyA-signals would be expected to predominate. Second, there is no reason to expect read-through rRNA transcripts to be spliced. Third, they appear to originate downstream of coding region, whereas read-through transcripts should be contiguous with the coding region. Indeed, all cell lines show a peak of CAGE tags in the general vicinity of the start of these transcripts. Neither the start of the transcripts nor the CAGE tag peaks fall in conserved regions, suggesting that either these transcripts are not conserved, the transcriptional start site does not need to be conserved at the sequence level, the conserved elements are too small to pass our threshold for a conserved block, or the conserved regulatory elements are located upstream or downstream of the TSS. The presence of transcripts, including some with the same splice junctions, in zone-1 in apes is further evidence that transcription in these regions may have functional significance. In contrast, zone-2 consistently shows chromatin marks associated with transcriptional activity across the cell lines we examined, but less consistent signals of actual transcripts. In addition, zone-2 lacks conserved IGS transcripts in any primate species we surveyed, suggesting that the conserved regions may not be associated with transcription. The pattern of conserved regions and open chromatin features in this zone suggest the conserved regions may have a function not associated with transcription. We suggest that enrichment of marks associated with active chromatin may be the result of these regions maintaining chromatin states that are important for rDNA function. Overall, given that the majority of IGS conserved regions fall into these zones and that the presence of active chromatin states has been documented in these regions previously [52,129], testing these zones for function is a high priority.

A major limitation of this and other studies looking at the rDNA is that the transcription and chromatin mapping results only give an average picture across all rDNA repeats, as mapping of sequence reads cannot currently distinguish individual rDNA repeats. Therefore, it is not possible to categorically associate factors such as chromatin marks of active transcription with transcripts, as the signals may come from physically distinct repeats. For example, there is evidence that some rDNA repeats exist outside of the nucleolus [130], and these may have a different transcriptional or chromatin profile to those located within the nucleolus. Similar limitations exist for trying to determine whether different histone modifications and transcription factors are located in the same rDNA repeats or not. Therefore, the chromatin profiles we observe might be an artificial composite of multiple, distinct states that exist in different rDNA units. Systems that are able to distinguish individual repeat units will be required to resolve these multi-copy issues of the rDNA.

The distinct nature of the embryonic cell line compared to the other cell lines is striking. This is most clearly seen in zone-2, where there are bi-directional small RNA peaks and a number of strong CAGE tag peaks that are specific to the stem cell line. Bi-directional small RNAs can act as enhancer RNAs [131,132], therefore it is possible that the bi-directional small RNA identified here is acting as a development-specific enhancer in embryonic stem cells [133]. rRNA transcriptional enhancers have been reported from*Xenopus*, *Drosophila*, mouse, and rat [134–138], but not human to date. Therefore, if this bi-directional small RNA is acting as an enhancer, it may be enhancing rRNA transcriptional activity. Evidence suggests that rRNA transcription is elevated in embryonic cell lines, and is downregulated to initiate differentiation [28–30,139]. Moreover, rRNA expression has been reported to be higher in certain embryonic cell lines than cancer cell lines [140]. Therefore, it will be interesting to determine whether this bi-directional small RNA plays any role in rRNA transcriptional regulation and pluripotency.

The rDNA units are arranged in loops inside the nucleolus [141], and this is facilitated by c-Myc [142]. This loop arrangement results from interactions between regions close to the rRNA promoter and terminators that are enriched for c-Myc [143], and interestingly these correspond to the promoter and zone-1, respectively. Recently, it has been shown that looping of rDNA units is also promoted by other regions of the IGS that interact with nucleolar matrix [144]. These regions correspond to conserved regions c15-c18, c31-c32, c33-c39, and c49, which also have c-Myc binding sites and many of which are enriched for c-Myc [144]. Interestingly, CTCF segmentation states that overlap c31-32 were predicted in zone-2 by Segway in all cell lines that had data. Based on our results and the roles that CTCF and c-Myc play in rRNA transcriptional regulation and genome organization [145,146], we speculate that some of the conserved regions play a role in mediating the threedimensional organization of the rDNA repeats in the nucleolus, facilitated by the association of CTCF and c-Myc with these regions [103,104,147].

In summary, our results provide a platform for comprehensively characterizing the functional landscape of the human IGS, and for developing a better understanding of the biological processes occurring in the rDNA and the nucleolus. They provide numerous predictions for functional elements in the IGS, in the form of conserved regions, and integrate a rich compendium of functional data to begin interpretation of the roles of these conserved regions. The strong association between the rDNA and human disease provides the impetus for characterizing functional elements in the IGS to better understand how they contribute to human health and wellbeing, and our results provide the basis from which to focus this functional characterization of the human rDNA.

## Acknowledgements

We thank Murray Cox (Massey University), Brian McStay (National University of Ireland, Galway), and the Ganley research group for helpful comments on the manuscript, and New Zealand Genomics Ltd for sequencing assistance. This work was supported by the Auckland Medical Research Foundation (4108008); the New Zealand Marsden Fund (14-MAU-053); and a PhD scholarship from the Institute of Natural and Mathematical Sciences, Massey University to SA.

## Supporting information

**S1 Appendix**. Sequence extracted from BAC clone GL000220.1 FASTA file.

**S2 Appendix**. Details of ChIP-seq, RNA-seq and CAGE data used in this study.

**S3 Appendix**. Additional supporting information from this study.

**S1 Supporting methods**. Additional details of methods used.

**Supporting_Figs.pdf: S1 Fig**. Estimating the lengths of rDNA units in primate BAC clones. **S2 Fig**. Repeat elements in the IGS of different primates species. **S3 Fig**. Sequence conservation of human rRNA transcriptional regulators. **S4 Fig**. Sequence conservation of potential c-Myc binding sites in the human IGS. **S5 Fig**. The transcriptomic landscape of the human IGS in different cell lines. **S6 Fig**. Chromatin, transcription factor and transcript landscape of the IGS in the umbilical vein endothelial cell line, HUVEC. **S7 Fig**. Chromatin, transcription factor and transcript landscape of the IGS in the lymphoblastoid cell line, GM12878. **S8 Fig**. Chromatin, transcription factor and transcript landscape of the IGS in the embryonic stem cell line, H1-hESC. **S9 Fig**. Chromatin, transcription factor and transcript landscape of the IGS in the hepatocellular carcinoma cell line, HepG2. **S10 Fig**. Chromatin, transcription factor and transcript landscape of the IGS in the leukemia cell line, K562. **S11 Fig**. Chromatin, transcription factor and transcript landscape of the IGS in the cervical carcinoma cell line, HeLa-S3. **S12 Fig**. Chromatin, transcription factor and transcript landscape of the IGS in the adenocarcinoma cell line, A549. **S13 Fig**. Genomic segmentation showing functional annotation states in the human IGS. **S14 Fig**. Transcripts in the Chimpanzee IGS. **S15 Fig**. Transcripts in the orangutan IGS. S16 Fig. Transcripts in rhesus macaque IGS. **S17 Fig**. Quantification of the expression level of IGS transcripts

**S1 Table**. Details of WGS data for the primates

**S2 Table**. Assembly statistics for the primate whole genome assemblies.

**S3 Table**. rDNA containing BAC clones identified by screening high-density BAC filters

**S4 Table**. rDNA sequence comparison between human and the six primate species.

**S5 Table**: The length variation between the WGA and BAC rDNA sequences of the six primate species.

**S6 Table**. Pairwise comparison between BAC clones rDNA sequences and WGA rDNA

**S7 Table**. Pairwise sequence comparisons showing the level of sequence conservation between human and ape Alu elements.

**S8 Table**. Details of conserved regions in the human IGS (base position corresponds to the human rDNA sequence extracted from BAC clone GL000220.1)

**S9 Table**. Sequence identity of the conserved regions with common marmoset and mouse rDNA.

**S10 Table**. Details of long poly(A+) IGS transcripts in HUVEC cell line (base position corresponds to the human rDNA sequence extracted from BAC clone GL000220.1).

**S11 Table**. Details of long poly(A+) IGS transcripts in GM12878 cell line (base position corresponds to the human rDNA sequence extracted from BAC clone GL000220.1).

**S12 Table**. Details of long poly(A+) IGS transcripts in the H1-hESC cell line (base position corresponds to the human rDNA sequence extracted from BAC clone GL000220.1).

**S13 Table**. Details of long poly(A+) IGS transcripts in HepG-2 cell line (base position corresponds to the human rDNA sequence extracted from BAC clone GL000220.1).

**S14 Table**. Details of long poly(A+) IGS transcripts in K562 cell line (base position corresponds to the human rDNA sequence extracted from BAC clone GL000220.1).

**S15 Table**. Details of long poly(A+) IGS transcripts in HeLa-S3 cell line (base position corresponds to the human rDNA sequence extracted from BAC clone GL000220.1).

**S16 Table**. Details of long poly(A-) IGS transcripts in the HUVEC cell line (base position corresponds to the human rDNA sequence extracted from BAC clone GL000220.1).

**S17 Table**. Details of long poly(A-) IGS transcripts in the GM12878 cell line (base position corresponds to the human rDNA sequence extracted from BAC clone GL000220.1).

**S18 Table**. Details of long poly(A-) IGS transcripts in the H1-hESC cell line (base position corresponds to the human rDNA sequence extracted from BAC clone GL000220.1).

**S19 Table**. Details of long poly(A-) IGS transcripts in the HepG-2 cell line (base position corresponds to the human rDNA sequence extracted from BAC clone GL000220.1).

**S20 Table**. Details of long poly(A-) IGS transcripts in the K562 cell line (base position corresponds to the human rDNA sequence extracted from BAC clone GL000220.1).

**S21 Table**. Details of long poly(A-) IGS transcripts in the HeLa-S3 cell line (base position corresponds to the human rDNA sequence extracted from BAC clone GL000220.1).

**S22 Table**. Details of the CAGE peaks in the human IGS in the selected cell lines (base position corresponds to the human rDNA sequence extracted from BAC clone GL000220.1).

**S23 Table**. Details of Long total RNA Chimpanzee IGS transcripts.

**S24 Table**. Details of poly(A+) Chimpanzee IGS transcripts.

**S25 Table**. Details of poly(A+) Orangutan IGS transcripts.

**S26 Table**. Details of poly(A+) RNA Rhesus macaque IGS transcripts.

